# Resource competition predicts assembly of *in vitro* gut bacterial communities

**DOI:** 10.1101/2022.05.30.494065

**Authors:** Po-Yi Ho, Taylor H. Nguyen, Juan M. Sanchez, Brian C. DeFelice, Kerwyn Casey Huang

## Abstract

Members of microbial communities interact via a plethora of mechanisms, including resource competition, cross-feeding, and pH modulation. However, the relative contributions of these mechanisms to community dynamics remain uncharacterized. Here, we develop a framework to distinguish the effects of resource competition from other interaction mechanisms by integrating data from growth measurements in spent media, synthetic community assembly, and metabolomics with consumer-resource models. When applied to human gut commensals, our framework revealed that resource competition alone could explain most pairwise interactions. The resource-competition landscape inferred from metabolomic profiles of individual species predicted assembly compositions, demonstrating that resource competition is a dominant driver of *in vitro* community assembly. Moreover, the identification and incorporation of interactions other than resource competition, including pH-mediated effects and cross-feeding, improved model predictions. Our work provides an experimental and modeling framework to characterize and quantify interspecies interactions *in vitro* that should advance mechanistically principled engineering of microbial communities.

## INTRODUCTION

Microbial communities are important for host health and environmental functions (Cho and Blaser, 2012; Singh et al., 2020), but their complex dynamics remain difficult to predict and engineer (Widder et al., 2016). A major challenge is that community members affect each other through a plethora of interaction mechanisms, including nutrient competition (Dal Bello et al., 2021; Hammarlund et al., 2021; Niehaus et al., 2019), metabolic cross-feeding (Adamowicz et al., 2018; Amarnath et al., 2021), pH modulation (Aranda-Díaz et al., 2020; Ratzke and Gore, 2018), toxins (Piccardi et al., 2019), and physical inhibition through secretion systems (Verster et al., 2017). The effects of many interaction mechanisms have been individually characterized in isolated contexts, but their relative contributions to the overall dynamics of a diverse community remain unclear in most cases.

Despite the complex nature of many natural microbiotas, community dynamics can often be captured by generalized Lotka-Volterra (gLV) models that describe interspecies interactions via phenomenological pairwise interaction coefficients (Faust and Raes, 2012; Fisher and Mehta, 2014; Venturelli et al., 2018; Xiao et al., 2017). These coefficients have often been inferred to be negative across a wide range of systems involving growth *in vitro* (Foster and Bell, 2012; Weiss et al., 2021), indicating that community members tend to inhibit one another. Competition for shared nutrients has been hypothesized to be a prevalent mechanism that generates negative interactions. Indeed, consumer-resource (CR) models in which interspecies interactions are mediated solely by resource competition can be mapped to gLV models with negative coefficients near steady state (Chesson, 1990; Cui et al., 2021; Xiao *et al*., 2017). Nonetheless, other mechanisms can play important roles in community assembly (Cordero and Datta, 2016; Venturelli *et al*., 2018; Weiss *et al*., 2021), and positive interactions can also be common in some situations (Kehe et al.). We therefore sought to clarify the origins of interactions by developing an integrated theoretical and experimental framework to disentangle the extent of resource competition from other interaction mechanisms.

To do so, we used CR models to integrate growth measurements in pairwise spent media (Biggs et al., 2017; Weiss *et al*., 2021), metabolomics (Han et al., 2021; Medlock et al., 2018), and 16S rRNA gene sequencing of assemblies of isolates. These techniques have been used previously to identify metabolites that mediate interspecies interactions (Biggs *et al*., 2017; Medlock *et al*., 2018), as well as to parametrize dynamical models for communities of a few species (Gowda et al., 2022; Hammarlund *et al*., 2021; Hart et al., 2019; Piccardi *et al*., 2019). Here, we extend these approaches to dissect interactions and measure the extent of resource competition in diverse communities and in contexts that mimic the complexity of natural environments like the mammalian gut.

We studied a collection of gut bacterial isolates that assembles into a community whose composition when grown *in vitro* can resemble the gut microbiota of mice colonized with the human fecal sample from which the isolates were obtained (Aranda-Díaz et al., 2022; Ng et al., 2019). We focused on 15 species that are representative of the phylogenetic diversity of human gut microbiotas (Fig. 1A), and together reconstitute ∼70% of the abundance of the parent community (Aranda-Díaz *et al*., 2022). We show that metabolomic profiles of the spent culture supernatants from each isolate can predict the amount of growth in pairwise spent media, and that these experimental data can be used to parametrize a coarse-grained CR model that accurately predicts community assembly. Similar qualitative conclusions were obtained in different growth conditions, suggesting that resource competition is generally a predominant factor driving *in vitro* community dynamics. Furthermore, we demonstrate a rational process to identify other interaction mechanisms, including cross-feeding and pH-mediated interactions, and to incorporate them into the model to improve predictions.

**Figure 1:**
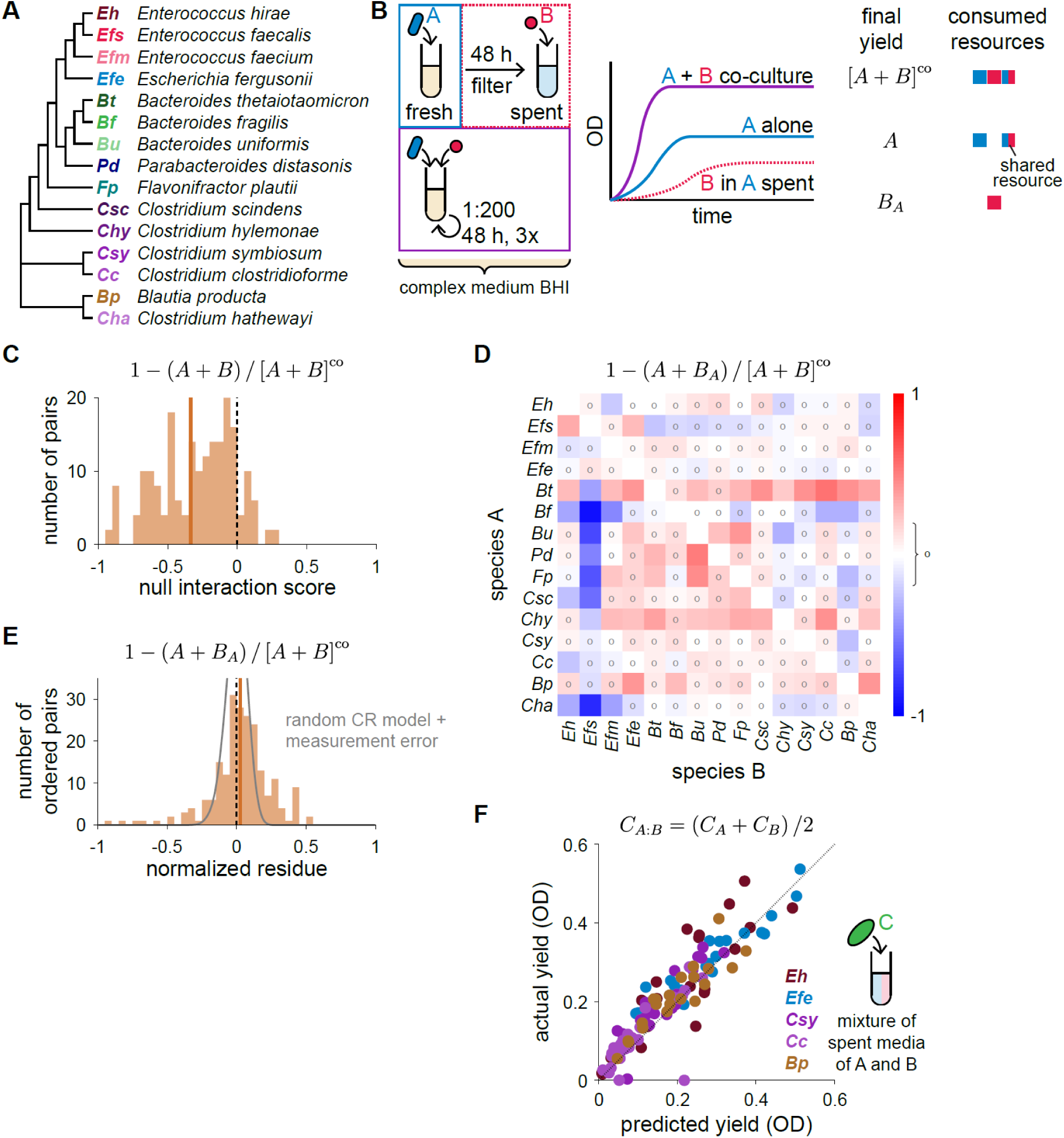
Coarse-grained resource competition describes most pairwise interactions. A) Phylogenetic tree of the 15 gut commensals studied here(Aranda-Díaz *et al*., 2022). The tree was constructed from the amplicon region of the 16S rRNA gene (Methods). B) Schematic of growth experiments in pairwise spent media and the predictions of the coarse-grained CR model. Growth curves of optical density (OD) over time were obtained for each species grown in isolation, in co-culture with every other species, and in the spent media of every other species, all in the complex medium Brain Heart Infusion (BHI). Experiments were replicated 2-4 times. In the coarse-grained CR model, the final yield is determined by the levels of coarse-grained resource groups, resulting in Eq. 1. C) The null interaction score, the difference between the yields of the co-culture and the sum of the isolate yields, was negative for most species pairs. Solid vertical line denotes the mean. D) Most components of the matrix of normalized resource competition residues are close to zero. Circles denote residues with absolute value <0.2. The null interaction scores and resource competition residues were calculated from mean values across 2-4 replicates. E) The distribution of normalized residues was centered about zero. Simulated results from randomly generated coarse-grained CR models with experimentally motivated measurement error are shown in gray (Methods). F) Yield in 1:1 mixtures of spent media is predicted by the average of the yield in each spent media individually. Colors denote the species grown, which was chosen to obtain a wide range of yields. Each of the species shown was grown in every pairwise mixture of the spent media from *Eh*, *Efe*, *Csy*, *Bt*, *Bp*, *Csc*, *Efs* or fresh BHI.

## RESULTS

### A coarse-grained CR model for inferring the origins of interspecies interactions

To characterize interspecies interactions in our model *in vitro* community (Fig. 1A), we measured the growth of each of the 15 species in isolation and in pairwise co-culture with each other species (Fig. 1B, Methods). All experiments were performed using the complex medium Brain Heart Infusion (BHI), which supported the growth of all isolates. In agreement with previous *in vitro* studies involving species from wide-ranging microbiotas, the gut commensals studied here typically inhibited the growth of one another in the sense that co-culture yields were less than the sum of the individual yields, as measured by optical density (Fig. 1C). Unless otherwise specified, all measurements were taken in stationary phase. Co-cultures were passaged until an ecological steady state was reached, in which the growth dynamics in successive passages were virtually identical (Methods). For each pair of species, we refer to the difference between co-culture yield and the sum of the two individual yields, normalized by the co-culture yield, as the null interaction score. The average null interaction score was −0.34 across the 105 pairs, out of which 93 exhibited negative scores (Fig. 1C). Although indicative of prevalent inhibition, negative null interaction scores cannot differentiate among the variety of inhibitory mechanisms that might be responsible.

Since resource competition is likely a common form of interspecies inhibition, we sought to quantify its extent by measuring the growth of each species in medium conditioned (hereafter “spent”, Methods) by the growth of each other species individually. Spent media exclude direct and physical effects that would emerge due to the presence of other species, but maintain environmentally mediated interactions, including resource competition, pH changes, toxins, and metabolic cross-feeding. To interpret the results, we considered a consumer-resource (CR) model in which resources are substitutable (i.e., any of the resources consumed by a species can support its growth), and are completely consumed and converted to biomass during growth to stationary phase (Erez et al., 2020; Ho et al., 2022). We coarse-grained the model by grouping metabolites that are consumed by the same set of species into one effective resource. While not a necessary assumption at this stage, coarse-graining simplified the interpretation of pairwise experiments and was important for analyzing assemblies of more than two species. A community of two species is then described by three effective resources: two specifically consumed by one of the two species, and one shared by both species. Metabolites that cannot be consumed by either species are ignored. With this coarse-graining, species A grown individually will consume its specific resource and the shared resource, leaving the other resource specific to species B in the spent medium, while all three resources will be consumed in a co-culture of the two species (Fig. 1B). Hence, if all species convert resources into biomass yield with the same efficiency, then the model predicts a simple relation linking the co-culture yield [𝐴 + 𝐵]^co^ to isolate growth in fresh and spent media,

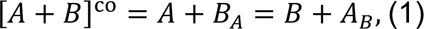

where 𝐴 and 𝐴_𝐵_ are the yields of A in isolation and in the medium spent by B, respectively, and similarly for 𝐵 and 𝐵_𝐴_. Small values of 𝑟(𝐴, 𝐵) = [𝐴 + 𝐵]^co^ − (𝐴 + 𝐵_𝐴_) and 𝑟(𝐵, 𝐴) = [𝐴 + 𝐵]^co^ − (𝐵 + 𝐴_𝐵_), which we refer to as resource competition residues, imply that the model can capture the assembly of the co-culture, presumably because interactions between A and B are dominated by resource competition. By contrast, large residues may highlight deviations due to interactions other than resource competition or differences in efficiency of resource conversion. Note that the possibility that species pairs can satisfy Eq. 1 while interacting via mechanisms other than resource competition cannot be ruled out, and that the two residues 𝑟(𝐴, 𝐵) and 𝑟(𝐵, 𝐴) for a pair of species can be asymmetric, potentially reflecting the directionality of interactions mediated by spent media. In addition to the assumptions above, Eq. 1 also ignores certain biological details, including saturation kinetics and hierarchical resource preferences. Nonetheless, Eq. 1 provides a useful baseline to interrogate the extent of resource competition as we demonstrate below.

### Most pairwise interactions can be described by resource competition

To quantify the relative contribution of resource competition to co-culture assembly, we applied Eq. 1 to interpret our pairwise spent media experiments. By contrast to the distribution of null interaction scores, the distribution of normalized resource competition residues 𝑟(𝐴, 𝐵)/[𝐴 + 𝐵]^co^ and 𝑟(𝐵, 𝐴)/[𝐴 + 𝐵]^co^ was centered about zero with mean 0.03 and standard deviation 0.21 across the 210 ordered pairs (Fig. 1D). Simulations of random instances of the CR models used to derive Eq. 1 (Methods) produced distributions of normalized residues centered around zero, as expected, and inclusion of 5% measurement noise, approximately equal to the standard error of the mean yields in our experimental data, broadened the distribution of residues to have maximum magnitude ∼0.2 (Fig. 1E, Methods). We therefore refer to a residue as near-zero if its magnitude is <0.2. Almost 3/4 of all residues (155/210) were near-zero and more than half of pairs (57/105) exhibited near-zero values for both normalized residues (Fig. 1D,E). Moreover, as a corollary to Eq. 1, the model predicts that the yield of species A in a 1:1 mixture of the spent media of B and C should be the average of the yields of A in each spent medium individually (Methods). This corollary was an excellent predictor of experimental data (Fig. 1F), suggesting that spent media typically did not exert other effects that would influence the consumption of nutrient mixtures. Taken together, these results demonstrate that Eq. 1 can describe most pairwise interactions, which suggests that in this community, resource competition is prominent compared with the contributions of other interaction mechanisms. Therefore, we will first focus on resource competition alone, and return to examine other mechanisms afterwards.

### Metabolomic profiles capture the landscape of resource competition

To further probe the nature of resource competition, we obtained untargeted metabolomics data via liquid chromatography coupled with tandem mass spectrometry (LC-MS) from the spent medium of each species (Fig. 2A, Methods). We detected thousands of features in fresh BHI, of which hundreds could be confidently annotated. The annotated metabolites included sugars, nucleotides, amino acids, and di- and tri- peptides, and represented diverse metabolic pathways (Han *et al*., 2021), suggesting that our pipeline provides a representative overview of bacterial metabolism. We therefore hypothesized that the spent media metabolomes reflect the landscape of resource competition, i.e., the extent of resource sharing among species (niches) as well as the approximate sizes of individual and shared niches. If true, then it should be possible to predict growth in spent media based on the metabolomic profiles.

**Figure 2:**
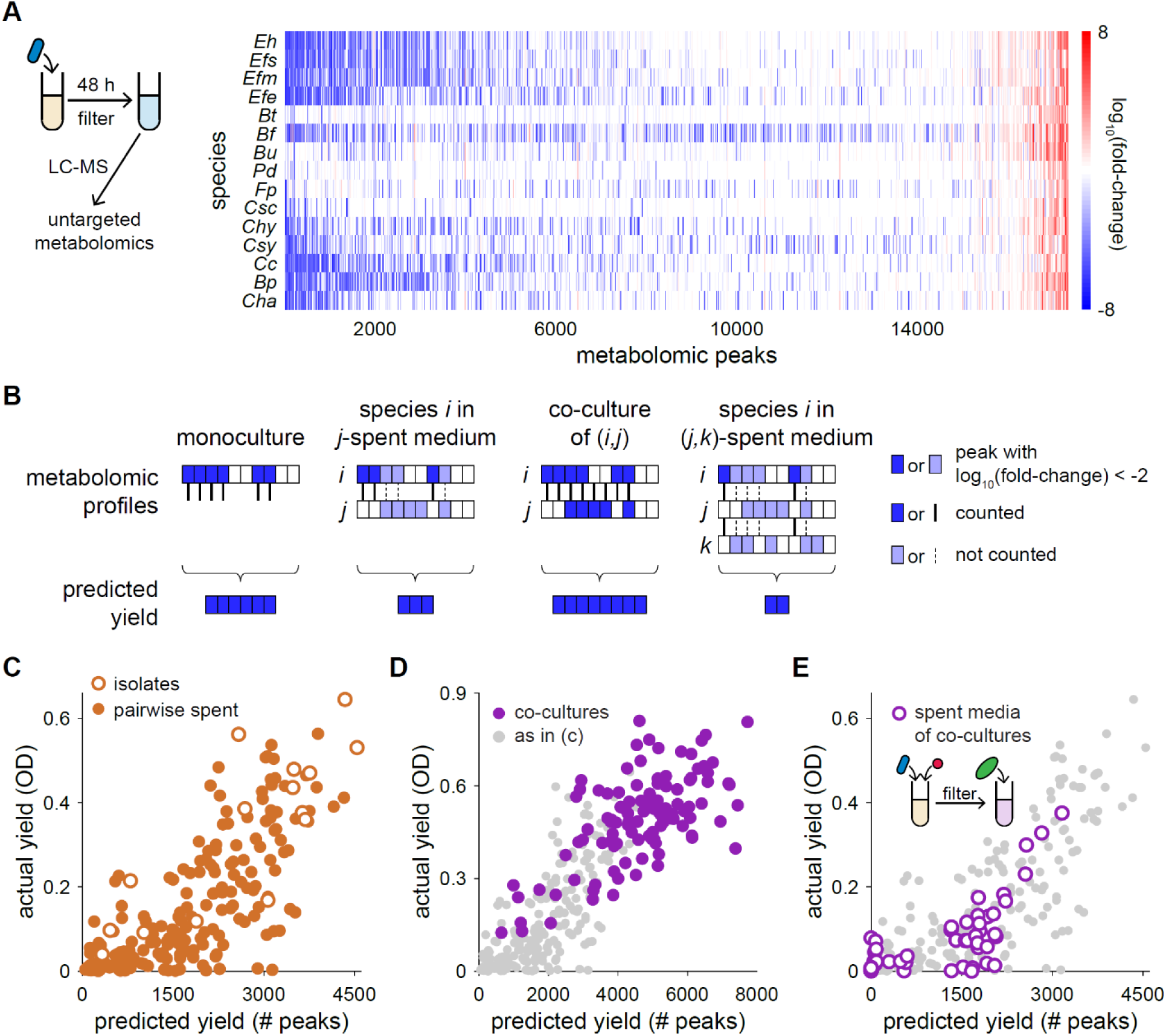
Metabolomic profiles predict growth in monoculture, co-culture, and spent media. A) Schematic of metabolomics experiments and the resulting profile of fold-change in LC-MS ionization intensity relative to fresh medium BHI for each species. Shown are all metabolomic features, including unannotated ones, that changed significantly in the spent medium of any of the species (Methods). For each metabolomic feature (“peak”), log_10_ of the average fold-change across 3 replicates is shown. One metabolite can generate multiple metabolomic features. B) Schematic of rule used to predict yield from metabolomics profiles. The predicted yield of a species in monoculture is defined as the number of metabolomic peaks (rectangles) that were depleted in the spent medium of that species (blue rectangles). The predicted yield of species 𝑖 in the spent medium of 𝑗 is defined as the number of peaks depleted by 𝑖 but not 𝑗, and analogously for co-cultures of 𝑖 and 𝑗, as well as 𝑖 growing in the co-culture of 𝑗 and 𝑘. C) Metabolomic profiles can generally predict yield measurements. The actual and predicted yields in monocultures and pairwise spent media experiments were highly correlated (Pearson’s correlation coefficient 𝜌 = 0.83, 0.76, and 0.78 for experiments involving isolates, pairwise spent media, and together, respectively). Shown are mean yields across 2-4 replicates. D) Metabolomic profiles successfully predict co-culture yields (𝜌 = 0.65). All co-culture experiments are shown. Shown are the mean yields across 2-4 replicates. Isolate and pairwise data, as in (b), are shown in gray in (c,d) as a visual guide. E) Metabolomic profiles successfully predict isolate yields when grown in the spent media of co-cultures (𝜌 = 0.74). Two species (*Eh* and *Efe*), chosen to obtain a wide range of yields, were grown in the spent media of all pairwise co-cultures of *Eh*, *Efe*, *Csy*, *Bt*, *Cc*, *Bp*, *Csc*, and *Efs*.

To connect metabolomes and growth measurements, we would ideally be able to relate the ionization intensities of a metabolite as reported by LC-MS to its contribution to biomass. However, one metabolite can generate multiple features (“peaks”) in LC-MS; moreover, the conversions from ion intensity to metabolite concentration can differ across metabolites (Alseekh et al., 2021; Han *et al*., 2021), and conversions from metabolite concentration to biomass can differ across species. In any case, these conversion factors are typically unknown. We reasoned that these details might be secondary to the total number of metabolites consumed in the limit of many involved metabolites, due to averaging over variations in these conversion factors. Accordingly, we tested the hypothesis that biomass is proportional to the number of peaks depleted, as defined by >100-fold depletion compared to fresh medium (Fig. 2B, Methods). This logic also predicts that the yield of species A in the spent medium of B should be proportional to the number of peaks depleted by A but not B. Since this hypothesis does not depend on the identity of the metabolites, unannotated features were also included to increase the number of metabolites and metabolic pathways involved. Deviations could be due to several causes, including biases in the conversions from peak height to concentrations, differences in the efficiency of biomass production across species, and interactions other than resource competition. Despite these potential limitations, the resulting predictions were highly correlated with experimental measurements of biomass yield (Pearson’s correlation coefficient 𝜌 = 0.78, Fig. 2C). Analogous predictions for co-cultures and growth in the spent media of co-cultures were also excellent (𝜌 = 0.65 and 0.74, respectively), and followed the same general trend as in the pairwise spent media experiments (Fig. 2D,E). Notably, successful predictions for growth in the spent media of co-cultures indicate that interactions among three species were also captured. These results further establish the importance of resource competition in this community, and demonstrate that metabolomic profiles can approximate the resource competition landscape.

### Resource competition predicts community assembly

Having found that resource competition can explain most pairwise interactions, we next sought to quantify the extent to which resource competition dictates the assembly of more diverse communities. The complex and undefined nature of many growth media such as BHI complicates efforts to directly manipulate resource levels. Instead, we reasoned that if resource competition is a leading driver of community assembly, then a CR model accounting for only resource competition without other interaction mechanisms should be able to predict community composition. We assembled subsets of varying sizes from the 15 species, passaged their mixture until they reached an ecological steady state as with co-cultures, and quantified community composition by 16S rRNA gene sequencing. We then parametrized a CR model by refining the approximate resource competition landscape provided by metabolomics using data from spent media experiments and tested whether it could predict community assembly (Fig. 3A, Methods).

**Figure 3:**
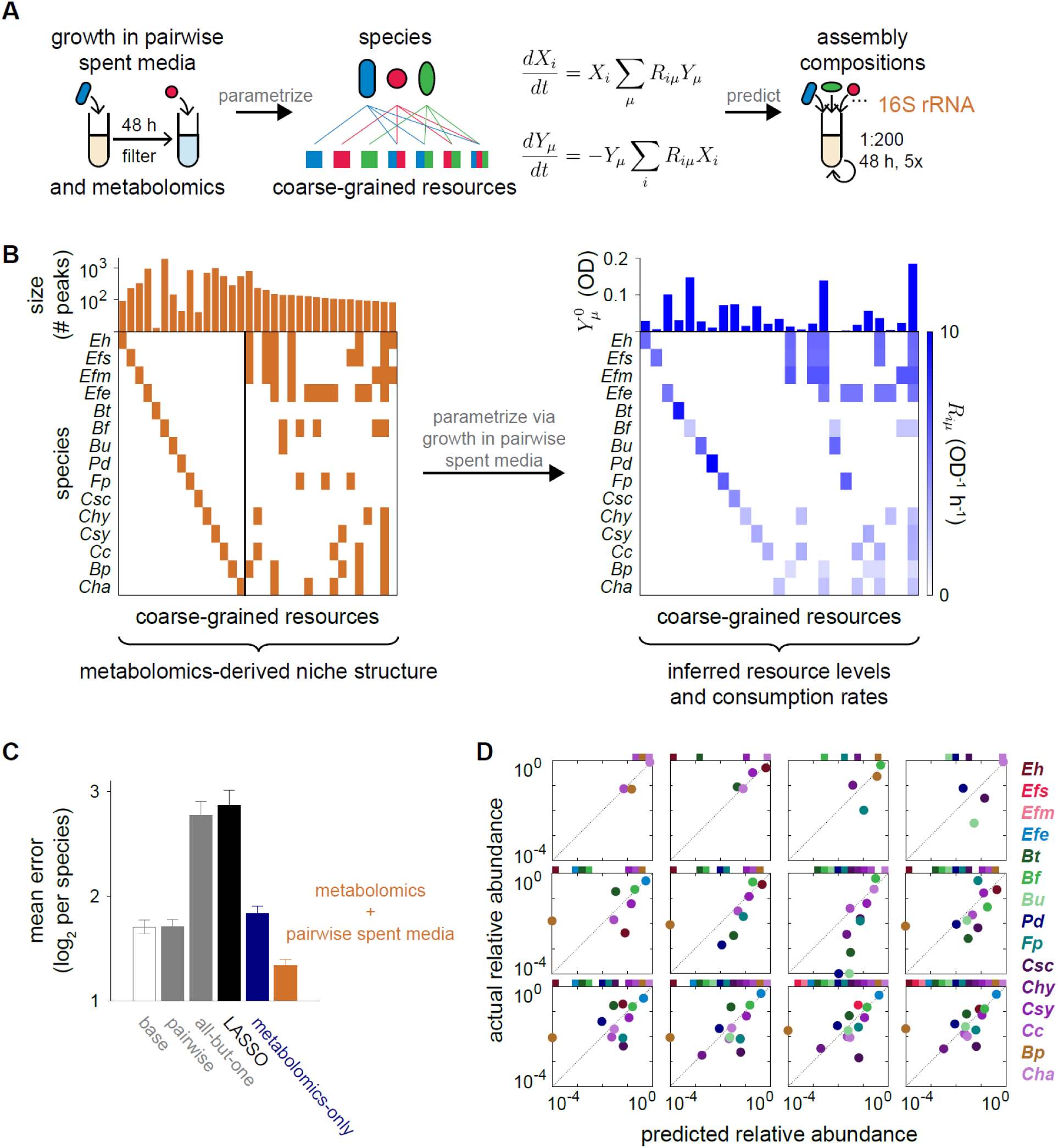
A consumer-resource model parametrized by metabolomics data and growth in spent media predicts community assembly. A) Schematic for predicting community assembly using a coarse-grained CR model. Metabolites are coarse-grained together if they are consumed by the same set of species (middle). The landscape of resource sharing among species was inferred from metabolomics data, and resource levels and consumption rates were inferred from growth curves in pairwise spent media (left). The parametrized CR model (Eq. 2, middle; implementation of lag times not shown for brevity) was then used to predict the composition of 185 assemblies of 2 to 15 species and compared against experimentally determined relative abundances from 16S rRNA gene sequencing (right) (Methods). B) The resource sharing structure obtained by coarse-graining metabolomics data in BHI (left). The 15 species-specific groups (left of vertical line) and 18 of the remaining resource groups with the greatest number of constituent metabolites (right of vertical line) are shown. The metabolomics-derived niche structure was used in combination with growth measurements in pairwise spent media to infer the resource levels 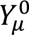 in fresh medium and resource consumption rates 𝑅𝑖𝜇 (right). C) A CR model parametrized by combining metabolomics and growth measurements in pairwise spent media achieved the best mean error out of all models considered. The mean error was determined by averaging across all 185 assemblies the magnitude of log_2_ fold-change between actual and predicted relative abundances, normalized by the number of species in the assembly. The performance of the metabolomics-derived landscape from (b) is shown in orange. The results of three hypothetical structures are shown in gray for comparison: a “base” structure with only species-specific niches, one including this base structure and every coarse-grained niche shared between species pairs, and another one including the base structure and every coarse-grained niche shared by all but one species. Shown in black and blue are the results of inferences via LASSO and the “metabolomics-only” scheme, respectively, as described in the text and Methods. D) Examples of predictions. Each panel represents one assembly, and colored squares, placed at the same location in each panel for each species, indicate species that were present in the inoculum of that assembly. The relative abundances of undetected species are set to 10^-4^ for visualization and for calculating prediction errors. Shown are mean values across 2-3 replicates.

Specifically, we considered the following CR model,

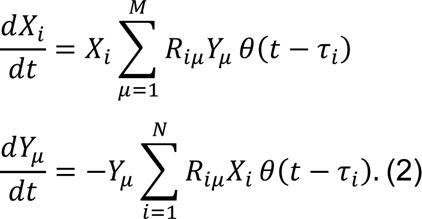

Here, 𝑋_𝑖_ denotes the abundance of species 𝑖, 𝑌_𝜇_ the amount of coarse-grained resource 𝜇, and 𝑅_𝑖𝜇_ the consumption rate of resource 𝜇 by species 𝑖. In addition, 𝜏_𝑖_ captures the lag time of species 𝑖, before which it does not grow nor consume resources. Lag time was implemented in Eq. 2 using a step function 𝜃, which is equal to zero if the input is <0 and equal to one otherwise. Lag times were estimated from growth measurements in monocultures and assumed to be the same in a community context as in monocultures (Methods). Eq. 2 explicitly describes the dynamics that gives rise to Eq. 1 in a community of two species, and like Eq. 1, ignores certain biological details including saturation kinetics and resource preferences. Nonetheless, the emergent behaviors of Eq. 2 have been previously shown to be able to quantitatively reproduce experimentally observed temporal fluctuations among wide-ranging microbiotas (Ho *et al*., 2022). We simulated Eq. 2 under serial dilution until species abundances reached an ecological steady state, mimicking our experimental protocol (Methods). The parameters of the model are the initial resource levels in fresh medium 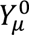 and the consumption rates 𝑅_𝑖𝜇_. For 𝑁 species, there are 𝑀 = 2^𝑁^ − 1 species combinations, and hence the same number of potential coarse-grained resources. Although coarse-graining removes some potential for niche partitioning by lumping together metabolites that could be consumed at relatively different rates by different species, it was important to simplify the parametrization process, and in hindsight, revealed the highly impactful shared niches that substantially affected community dynamics. Under coarse-graining, the challenge is to infer the 2^𝑁^ − 1 resource levels and 𝑁 × (2^𝑁^ − 1) consumption rates from the far fewer 𝑁^2^ growth curves in spent media and 𝑁 metabolomic profiles of the isolates. We decomposed the challenge into three steps (Methods).

First, the subset of coarse-grained niches that have non-zero resource levels 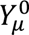, or equivalently, the subset of consumption rates 𝑅_𝑖𝜇_ that are non-zero, was determined from metabolomics data via coarse-graining as detailed below. Second, given this structure of resource consumption, the corresponding resource levels in fresh medium 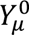 were inferred from the experimentally determined yield of species 𝑖 in the spent medium of 𝑗, which the model predicts to be 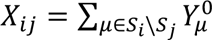 where 𝑆_𝑖_ is the set of resources consumed by species 𝑖, and \ denotes the difference between sets – in other words, the sum is over resources 𝜇 consumed by 𝑖 but not 𝑗 such that 𝑅_𝑖𝜇_ > 0 but 𝑅_𝑗𝜇_ = 0. If the number 𝑀 of coarse-grained resources under consideration is less than the number of measurements 𝑁^2^, this problem is constrained and can be best fit by linear regression. Finally, the consumption rates 𝑅_𝑖𝜇_ were inferred from experimentally determined growth rates, which the model predicts to have a maximum value of 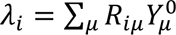 for species 𝑖 in monoculture. Although a linear regression similar to the one for resource levels can be carried out using the growth rates in spent media to determine 𝑅_𝑖𝜇_, we decided to simplify the problem given limitations in the accuracy of growth rate measurements in cultures with low yield. We assumed that 𝑅_𝑖𝜇_ = 𝑅^∗^ for all resources 𝜇, i.e., species 𝑖 consumes all resources that it uses at the same rate (e.g., (Good et al., 2018; Tikhonov and Monasson, 2017)), and hence, 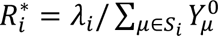 .

Since the latter two steps are simpler to solve, the core challenge is to choose the subset of consumption rates 𝑅_𝑖𝜇_ that are non-zero, i.e., the resource utilization structure. Crucially, metabolomics data can directly reveal the potential niche overlaps among multiple species, as demonstrated by their ability to predict growth in spent media (Fig. 2). We therefore used the metabolomics data to guide our choice of the utilization structure (Fig. 3B). The >15,000 features that were depleted in at least one spent medium were grouped into 1,211 coarse-grained resource niches (Fig. S1A). Most metabolites clustered into large groups: the 100 groups with the largest number of constituent metabolites comprised ∼84% of the metabolites (Fig. S1B). Each species was associated with a set of metabolites that it uniquely depleted, which collectively comprised ∼49% of the metabolites (Fig. S1A). To predict community compositions from these metabolomics-derived niches, we first restricted our analysis to the resource utilization structure determined by the 15 species-specific niches and a subset of the remaining niches with the largest number of constituents, reasoning that they should encode most of the information about the landscape of resource competition since the number of depleted peaks was strongly correlated with biomass yield (Fig. 2). We varied the number of niches included and compared model predictions against 185 assemblies of 2 to 15 species. Mean model error was minimized for the structure containing the species-specific niches and 18 of the largest remaining niches (Fig. 3C), which together comprised ∼68% of the metabolites. Remarkably, the inferred resource levels and consumption rates (Fig. 3B, S2A) predicted assembly compositions with a mean error of 1.33 log_2_ fold-change per species (“doublings per species”; defined as 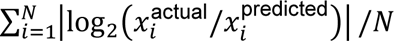, where 𝑥_𝑖_ is the relative abundance of species 𝑖, which was set to 10^-4^ if species 𝑖 was undetectable), which was smaller than the error of other models considered, as discussed below (Fig. 3C,D, S2B, Methods). This analysis ignores potential biases that can arise from conversions between biomass yields, optical density measurements, and 16S counts that remain challenging to quantify. Nonetheless, the close agreement between the CR model and experimental observations suggests that resource competition drives community assembly to a large extent in our system.

### Properties of resource competition in our community

To further evaluate the relevance of metabolomics-derived niches, we attempted to predict community compositions using hypothetical structures of resource consumption. First, we tested a “base” structure that includes only species-specific resources. Next, we added to the base structure either every coarse-grained resource shared only between species pairs, or every resource shared across all but one species (Fig. S3A). All three structures predicted community compositions less accurately (mean error of 1.70, 1.71, and 2.77 doublings per species, respectively; Fig. 3C) than the CR model including all species-specific niches and the next 18 largest niches, indicating that the hypothetical structures failed to fully capture the landscape of resource competition in our community. In fact, the set of all-but-one niches significantly decreased model performance relative to the base structure. Thus, the incorporation of low-relevance niches can decrease model performance despite providing additional degrees of freedom.

Having established the relevance of metabolomics-derived niches, we next investigated their implications for the nature of resource competition in our community. First, the presence of species-specific niches for all 15 species explained their widespread coexistence in assemblies because in the absence of other antagonistic interactions, each species can grow by accessing its exclusive set of metabolites. Given only the species-specific niches, our modeling framework would infer the species-specific resource levels, and hence the species abundances in community, to be proportional to the average yield across monoculture and pairwise spent media for each species. The ensuing base model predicted assembly compositions to a reasonable degree, although 28% less accurately than the best model (Fig. 3C). This result mechanistically explains a previous finding that isolate yield correlates with abundance in a community context (Aranda-Díaz *et al*., 2022). In addition to the species-specific niches, there was also substantial resource sharing among groups of species (Fig. 3B). The shared metabolite groups comprise approximately half of all metabolomic features that were depleted, and within the inferred model, accounted for approximately half of the total resource levels (Fig. 3B). Inclusion of shared resources improved model predictions, but prediction error was non-monotonic with respect to the number of niches included (Fig. S3B), highlighting the combinatorial complexity of the resource competition landscape.

A key assumption in our model is that the metabolic functions of individual species remain unchanged in the context of a community. Several lines of evidence support this assumption. First, the compositions of the full 15-member community and dropout assemblies with 14 of the 15 species were well predicted by the CR model with parameters inferred from mono- and co-cultures (mean error of 1.63 doublings per species, Fig. S2B), suggesting that the metabolic capacities of the species remained largely unchanged even in the most complex assemblies possible with the species considered here. Mixing, or “refilling”, dropout assemblies with the species that was left out mostly resulted in communities with compositions indistinguishable from the full community assembled from monocultures (Methods, Fig. S4), in agreement with our CR model, which predicts that initial abundances do not affect steady-state values. Taken together, these results imply that community assembly can be accurately represented by a CR model with fixed parameters regardless of which species are present.

### Comparisons with other models

To test other inference approaches, we determined the landscape of resource competition based on all >1,000 metabolomics-derived coarse-grained resources via regularized regression, i.e., LASSO, against yields in pairwise spent media (Methods). The resulting predictions for assembly compositions were substantially worse than even the base structure (mean error of 2.87 doublings per species, Fig. 3C), again highlighting that the incorporation of low-relevance niches can decrease model performance. We then tested a simple scheme in which the relative abundance of a species in a community was defined to be the fraction of metabolites that it consumed out of the union of metabolites consumed by all species in the community, and when multiple species shared the same set of metabolites, each species was assigned a fraction of those metabolites in proportion to its monoculture yield. Predictions based on this simple “metabolomics-only” scheme were also less accurate than for the base structure (mean error of 1.84 doublings per species, Fig. 3C), further strengthening the utility of our framework that combines metabolomics and growth measurements for accurate predictions.

To provide a comparison for the various CR models, we used the pairwise spent media experiments to parametrize a gLV model with pairwise interactions (Methods). The resulting gLV model failed to accurately predict assembly compositions, resulting in a mean error of 2.48 doublings per species, notably with a mean error of 3.98 doublings per species in assemblies of more than two species. One obvious cause of the disagreements was the prevalence of negative interactions, which led the gLV model to predict extinctions in cases that were not observed experimentally (Fig. S5). The relatively poor performance of other models further supports our conclusion that resource competition is a major driver in our system, and validates our method to parametrize a predictive CR model. Taken together, these results demonstrate that our framework, despite its potential limitations, led to the best parametrization of the landscape of resource competition that we have identified thus far.

### The prominence of resource competition across environments

Having developed an experimental and theoretical framework to probe resource competition, we next sought to examine the generality of the prominence of resource competition across media. To do so, we applied the framework developed in BHI to the same 15 species grown in another commonly used complex medium, modified Gifu Anaerobic Medium (mGAM). mGAM shares some but not all ingredients with BHI, and as a result, many species grew differently in the two media (Fig. 4A). In particular, the four species in the Bacteroidetes phylum (*Bacteroides thetaiotaomicron*, *Bacteroides fragilis*, *Bacteroides uniformis*, and *Parabacteroides distasonis*) exhibited substantially larger yields in mGAM. Despite these differences, spent media experiments showed that the distribution of normalized resource competition residues in mGAM was centered about zero (Fig. 4B), suggesting that resource competition is prominent in mGAM as it is in BHI.

**Figure 4:**
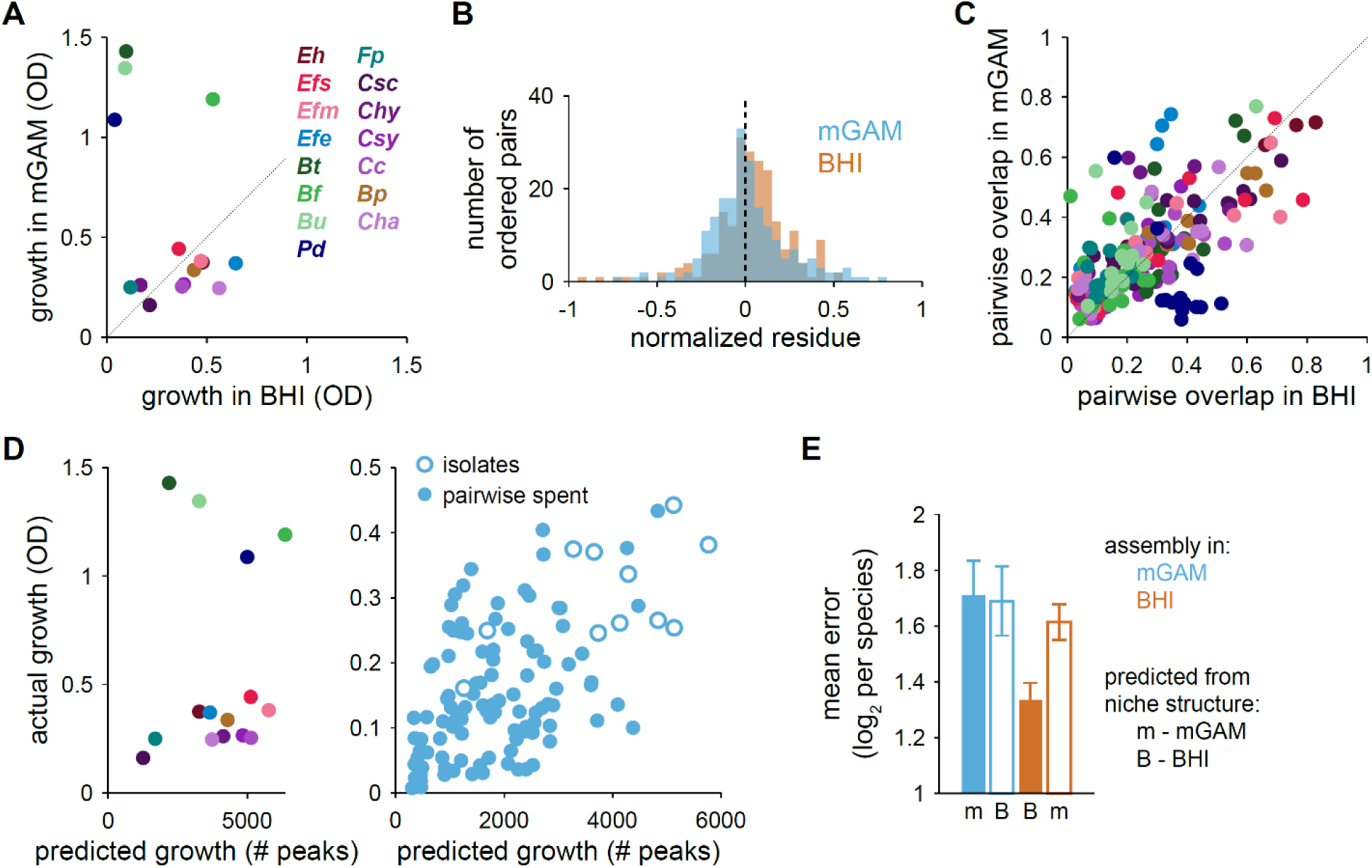
Resource competition drives community assembly across distinct growth environments. A) Monoculture yields in mGAM differed from those in BHI, particularly for the Bacteroidetes, which exhibited substantially larger yields in mGAM. Mean values across 2-4 replicates are shown. B) The distribution of resource competition residues in mGAM was centered about zero as in BHI. C) Pairwise overlaps in metabolomic profiles in mGAM and BHI were moderately correlated (𝜌 = 0.66). The pairwise overlap between the ordered species pair (𝑖, 𝑗) is defined as the number of metabolomic peaks depleted by both species divided by the number depleted by species 𝑖. Pairwise overlaps are colored according to species 𝑖. D) The experimentally determined yield in monoculture (left) or pairwise spent media (right) was correlated with the number of depleted peaks for experiments not involving the four Bacteroidetes species (𝜌 = 0.54), which are not shown on the right. E) Mean error of model predictions in mGAM (blue) and BHI (orange), as inferred using resource utilization structures derived from metabolomics data in mGAM (“m”) or BHI (“B”), with the resource levels and consumption rates re-inferred. Mean errors resulting from utilization structures derived from metabolomics data in the matching environment are shown as solid bars, while mean errors from mis-matched structures are shown as empty bars.

Following the same logic as in BHI, we obtained LC/MS measurements of the spent media of each of the 15 isolates grown in mGAM. The resulting sizes of pairwise niche overlaps were distinct but correlated across the two media (Fig. 4C), indicating that the landscape of resource competition depended partially on the environment. In addition, the mapping from metabolomes to growth yield was distinct across the two media. The number of depleted peaks in mGAM was correlated with growth in spent media (𝜌 = 0.54, Fig. 4D), although not as strongly as in BHI and not for the Bacteroidetes (Fig. 4D). This result suggests that the substantial increase in yield for the Bacteroidetes was due to an increase in the efficiency of biomass generation in mGAM, for example due to the presence of cofactors like hemin and vitamins (Halpern and Gruss, 2015).

Despite potential differences in the efficiency of biomass generation for the Bacteroidetes, we reasoned that metabolomics data should still reflect the utilization structure of the shared resources. When parametrized using an optimized set of the largest metabolomics-derived niches, our CR model predicted assembly compositions in mGAM with a mean error of 1.72 doublings per species (Fig. 4E). Although these predictions were slightly less accurate than in BHI, they were still more accurate than other models that we considered. Predictions of assembly in BHI were less accurate when using the mis-matched mGAM-derived niches (with the resource levels and consumption rates re-inferred based on the mGAM-derived utilization structure) versus the matching BHI-derived niches (Fig. 4E), in agreement with the observation (Fig. 4C) that the landscape of resource competition can depend to a degree on the environment while also being partially fixed by the identity of the species. By contrast, the accuracy of predictions in mGAM were not significantly affected when using mis-matched niches (Fig. 4E). This finding suggests that our framework did not capture certain mGAM-specific interactions, consistent with the weaker correlation between metabolomes and yields in mGAM (Fig. 4D). Nonetheless, our CR modeling framework predicted assembly compositions in mGAM with similar efficacy as in BHI. Taken together, these findings establish the prominence of resource competition across two distinct environments.

### A framework to disentangle interaction mechanisms

While most pairs of species exhibited resource competition residues that were near-zero, ∼25% of the residues in BHI deviated from zero (Fig. 1D), suggesting that some species additionally interact via mechanisms other than resource competition. Deviations from Eq. 1 can arise in many ways. For example, if the growth of species A affects that of B by an amount Δ in addition to the assumptions of resource competition underlying Eq. 1 and this effect occurs similarly in spent medium and in co-culture, then the model would predict that 𝑟(𝐴, 𝐵) ≔ [𝐴 + 𝐵]^co^ − (𝐴 + 𝐵_𝐴_) = 0 and 𝑟(𝐵, 𝐴) ≔ [𝐴 + 𝐵]^co^ − (𝐵 + 𝐴_𝐵_) = Δ (Fig. 5A). If the effect of A on B is specific to spent medium and does not occur in co-culture, the model would instead predict 𝑟(𝐴, 𝐵) = −Δ and 𝑟(𝐵, 𝐴) = 0 (Fig. 5A). For example, a species involved in the latter scenario is *Blautia producta* (*Bp*), whose spent medium almost completely inhibited the growth of all other species, i.e., Δ < 0 (Fig. 5B). However, *Bp* grew more slowly than many other species (Fig. S2A), and thus these other species were able to grow in co-culture before the inhibitory effects of *Bp* occurred. The residues 𝑟(*Bp*, 𝐵) were therefore >0 for most other species B (Fig. 5C), implying that there is a surplus of growth in co-culture relative to the inhibitory effects of *Bp*-spent medium. *Bp*-spent medium was highly acidic (pH ≈ 5), and the growth inhibition of other species was largely lifted in *Bp*-spent medium that was adjusted to neutral pH (Fig. 5B, Methods). Importantly, residues computed from yields in the neutralized spent medium were less positive and closer to zero (Fig. 5C), demonstrating that pH neutralization brought these pairs into close agreement with the baseline CR model and Eq. 1. Thus, the combination of resource competition and pH modification can describe the interspecies interactions of *Bp*.

**Figure 5:**
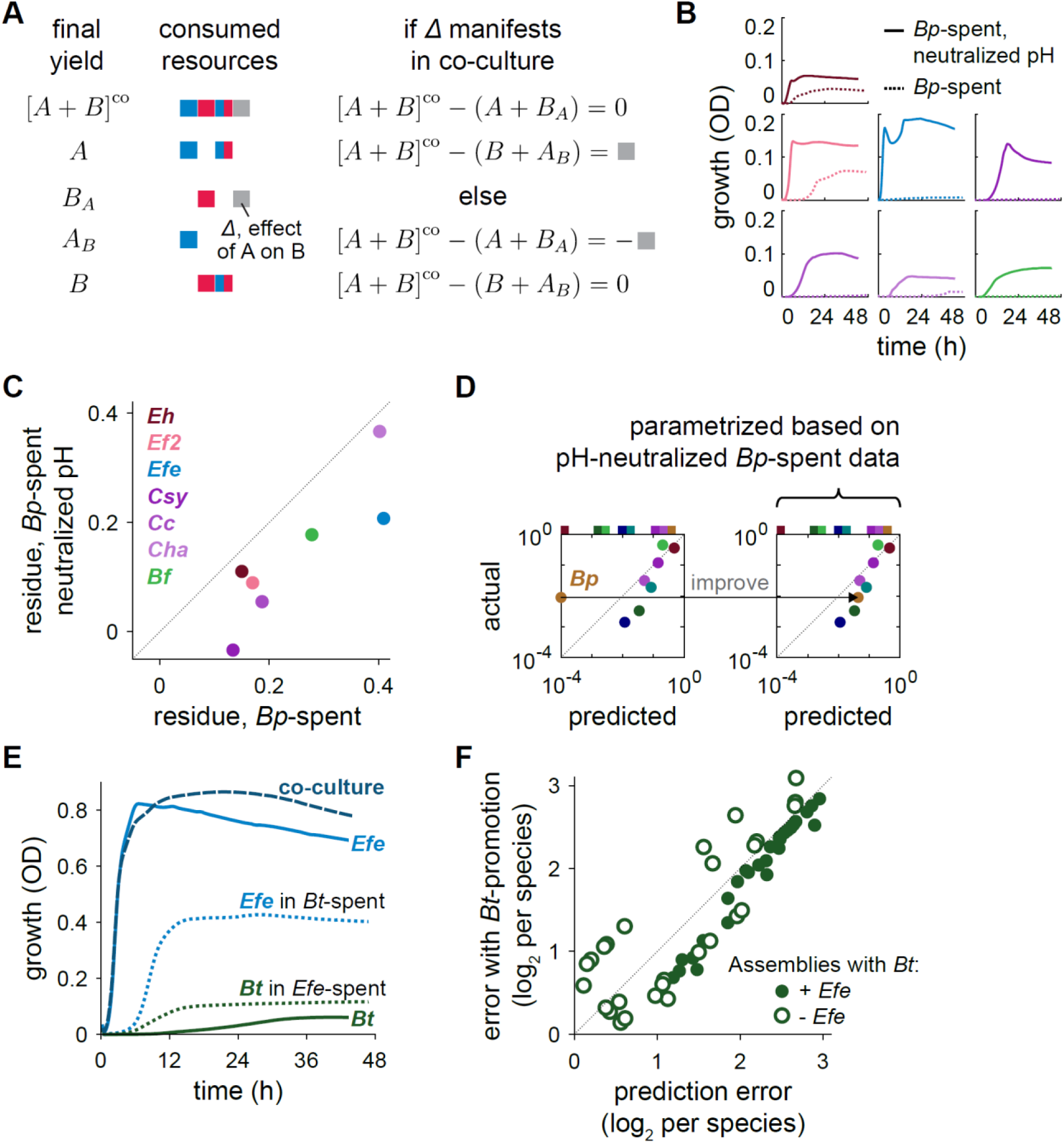
Strategies for disentangling pH and metabolic cross-feeding interactions from resource competition. A) Schematic for interpreting resource competition residues. Gray square denotes an additional contribution to growth from a mechanism other than resource competition. B) pH-mediated interactions involving *Blautia producta* (*Bp*). Shown are growth curves in *Bp*-spent medium and *Bp*-spent medium with neutralized pH for species that grow more quickly than *Bp* in monoculture. C) Resource competition residues became less positive and closer to zero after neutralizing the pH of *Bp*-spent medium (Methods). D) Model predictions improved after parametrization based on growth in pH-neutralized *Bp-*spent medium. Shown are predictions for an example assembly from Fig. 3D based on models parametrized without (left) and with (right) data from pH-neutralized *Bp*-spent medium. Arrow highlights improved model prediction for *Bp* coexistence. E) *E. fergusonii* interacts with *B. thetaiotaomicron* through cross-feeding. Shown are OD over time in pairwise spent media for *Ef*e and *Bt*. *Bt* grew more quickly and to a higher yield in *Efe*-spent medium, the only case of growth promotion in spent medium out of all 210 pairs. F) Errors of model predictions after incorporating cross-feeding into the model. Prediction errors for each assembly containing *Bt* are shown for the parametrized CR model (Fig. 3B) and the same model with an additional boost in *Bt* growth. A fixed boost to *Bt* abundance equal to the difference in yields between *Bt* in *Efe*- spent and *Bt* in monoculture was applied whenever *Bt* was present. Assemblies that also contain *Efe* are shown as filled circles, and those without *Efe* are shown as empty circles. Assemblies with *Efe* were always better predicted when *Bt*- promotion was included, whereas assemblies without *Efe* fared better or worse in an apparently random manner.

Within a model that accounts for only resource competition, growth inhibition can only be due to niche overlap. Therefore, the outsized inhibition by *Bp*-spent medium caused the linear regression used for parametrizing resource levels to infer high levels for niches shared between *Bp* and other species but a zero level for the *Bp*-specific niche (Fig. 3B). As a result, *Bp* was often predicted to go extinct (Fig. 3D), in disagreement with experimental data (mean error in *Bp* relative abundance of 4.07 doublings, largest out of all species). By contrast, when yields from pH-neutralized *Bp*-spent medium were used to parametrize the CR model, the *Bp*-specific niche was inferred to have a non-zero resource level, which dramatically improved model predictions (mean error in *Bp* abundance of 1.58 doublings, mean error across all species of 1.31 doublings per species, Fig. 5D). These findings highlight that while mechanisms other than resource competition can potentially confound model parametrization, their effects can be disentangled and model parametrization improved in a quantitative and rational manner.

Metabolic cross-feeding is another potential interaction mechanism that can occur in addition to competition for existing resources. Of the >17,000 metabolomic features in BHI that changed significantly in the spent media of any of the species, <2,500 (<15%) were produced (increased by >10-fold relative to fresh medium) by at least one species. Of these produced metabolites, <800 (<5%) were consumed by at least one other species (Fig. 2A). This analysis does not account for the ∼700 peaks (<5%) that had an intensity <10^2^ in fresh medium, which would make their consumption undetectable based on our definition. Nonetheless, the low percentages of produced and cross-feeding metabolites detected suggest that cross-feeding interactions are uncommon or exert small effects in our community. Substantial growth promotion by spent media was indeed rare. Only a single ordered pair of species out of 210 exhibited strong enough growth promotion such that growth in spent medium surpassed that in fresh medium: the spent medium of *Escherichia fergusonii* (*Efe*) substantially boosted the growth of *Bacteroides thetaiotaomicron* (*Bt*), resulting in a positive residue, 𝑟(*Bt, Efe*) > 0 (Fig. 5E). *E. coli* (a close relative of *Efe*) can promote the growth of *Bt* due to the production of porphyrins, cofactors involved in iron metabolism that can stimulate the growth of members of the Bacteroidetes phylum (Halpern and Gruss, 2015). The yield of *Bt* grown in BHI supplemented with hemin, which contains porphyrins, increased to a similar extent as in *Efe*-spent medium (Fig. S6A), suggesting that the same cross-feeding mechanism occurs between *Efe* and *Bt* as between *E. coli* and *Bt*. Moreover, mGAM contains hemin, and *Bt* grew substantially better in mGAM than in BHI (Fig. 4A). Importantly, the residues between *Efe* and *Bt* were near-zero in mGAM, supporting the notion that interactions can depend on the environment.

The beneficial effects of *Efe* on *Bt* persisted in a community context. In particular, *Bt* was not detected in the dropout assembly in which *Efe* was removed. To systematically quantify the effects of removing one species on all other species, we calculated *z*-scores 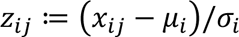, where 𝑥_𝑖𝑗_is the log_10_ relative abundance of species 𝑖 in the dropout assembly in which species 𝑗 was removed, and 𝜇_𝑖_ and 𝜎_𝑖_ are the mean and standard deviation, respectively, of the log_10_ relative abundance of species 𝑖 across all dropout assemblies. Of all *z*-scores, only *Bt* and *Parabacteroides distasonis* (*Pd*) in the *Efe*- dropout had absolute value >3 (Fig. S6B), indicating significant interactions. Although the yield of *Pd* did not increase in *Efe*-spent medium, its growth rate increased (Fig. S6C), corroborating the beneficial effects of *Efe* on *Pd* suggested by their significant *z*-score. The rarity of significant *z*-scores was consistent with the rarity of substantial growth promotion in spent media. More broadly, these results indicate that pairwise interactions other than resource competition can persist in larger communities.

The parametrization of the CR model used above to predict assemblies in BHI did not incorporate the beneficial effects of *Efe* on *Bt*, which we hypothesized would cause poor predictions for the relative abundance of *Bt* when *Efe* was present. To test whether incorporating this beneficial interaction improves predictions, we modified the CR model by assuming that whenever *Efe* and *Bt* were both present, the predicted abundance of *Bt* increases by a constant amount equal to the difference in yields between *Bt* grown in *Efe*- spent medium and in fresh BHI. Remarkably, without any additional tuning of model parameters, prediction errors decreased for all assemblies containing both *Efe* and *Bt* (Fig. 5F). By contrast, when the same increase in abundance was applied to *Bt* when *Efe* was absent, prediction errors increased in some cases (Fig. 5F), implying that the enhanced growth of *Bt* was *Efe*-dependent. This example demonstrates that metabolic cross-feeding can be quantitatively and straightforwardly incorporated into the CR model.

## DISCUSSION

In this study, we developed an experimental and theoretical framework to quantify the contributions of resource competition to community assembly. Although our framework cannot yet distinguish among all possible mechanisms, its quantification of resource competition led to accurate predictions of diverse assemblies. Similarly accurate predictions were obtained across two complex media, suggesting that the predominance of resource competition can be general across environments. Importantly, we identified and quantified several other interaction mechanisms to improve model predictions. These results provide a broadly applicable null model for community dynamics *in vitro* and *in vivo*.

For example, our findings unify observations from several previous studies involving *in vitro* communities. For the same model community studied here (Aranda-Díaz *et al*., 2022), the effects of the addition of simple carbon sources on community dynamics could be quantitatively explained by the monoculture growth behaviors of isolates on those carbon sources, implying that competition for resources again drove community dynamics. When a separate synthetic community of >100 gut commensals colonizing germ-free mice was challenged by a human fecal sample via gavage, the relative abundances of species that persisted post-challenge were highly correlated with their pre-challenge values (Cheng et al., 2021). This observation mirrors the tight distribution of *z*-scores in dropout assemblies and the compositional similarity of refilled communities, which together show that the removal or addition of a species typically did not affect community composition. This correspondence suggests that resource competition is predominant even in a community almost an order of magnitude more diverse than the one we have studied, and even in the context of host colonization.

Our model should be able to predict the outcomes of *in vitro* scenarios such as nutrient perturbation, resistance to invasion, and community coalescence, which have direct implications for the *in vivo* analogs of dietary switches, pathogen infection, and fecal microbiota transplantation, respectively. Microbiota-accessible carbohydrates like inulin simultaneously affect community composition and decrease burden from *C. difficile* infection in mouse models (Hryckowian et al., 2018). Decrease in *C. difficile* burden was linked with short chain fatty acids, metabolites associated with microbial metabolism of complex carbohydrates whose production by *Bacteroides* species has also been implicated in colonization resistance against *Salmonella* (Jacobson et al., 2018). The interplay among diet, community composition, and colonization resistance can be further clarified by measuring resource competition landscapes in media supplemented with complex carbohydrates. Model predictions from the resulting niche overlaps can untangle metabolite- or host-mediated effects from resource competition. Conversely, therapy by fecal microbiota transplantation seeks reliable colonization, the extent of which can also be predicted from *in vitro* growth measurements.

A key conclusion that emerges from our study is that complexity can ultimately generate simplicity. In fact, the diversity of a community likely contributes to the predominance of resource competition by dampening the effects of outlier interactions on the rest of the community. For example, although *E. fergusonii* substantially promoted the growth of *B. thetaiotaomicron* (Fig. 5E), this interaction was the only case of growth promotion in spent media out of 210 ordered pairs and thus did not significantly affect other species in a community context. By contrast, the fly gut microbiota consists of only five species, hence the cross-feeding and pH interactions observed among those species can strongly affect the overall dynamics of the community (Aranda-Díaz *et al*., 2020).

The complexity of the environment also likely contributes to the predominance of resource competition. In particular, a complex medium may provide metabolites that would otherwise be cross-fed in minimal environments. Therefore, the environment may be as important as the identity of the community members in determining community dynamics (Hart *et al*., 2019; Momeni et al., 2017). The complexity of the environment also likely contributes to the simple mapping between the number of metabolites and biomass yield (Fig. 2), presumably by averaging over biomass contributions from numerous sources. Consequently, a relatively simple resource competition landscape emerged (Fig. 3b). The media studied here were more complex than the community in the sense that they provided exclusive niches for each species (Fig. 3B), thereby enabling widespread coexistence. Since the complexity of the environment relative to that of the community is an important determinant of the behavior of CR models (Cui *et al*., 2021), it will be insightful to investigate the resource competition landscape in more sparse, ideally defined environments for which not all members have species-specific niches.

The predominance of resource competition enabled our model to capture the majority of interactions, and hence, predict community assembly to reasonable accuracy despite initially not accounting for other interaction mechanisms. Some interactions likely only manifest in pairwise spent media and do not affect the dynamics of larger communities. For example, although pH modification by *Bp* confounded model parametrization (Fig. 5B-D), it evidently played a relatively unimportant role in community dynamics, likely because most other species grew before *Bp* could modify the environment. Another phenomenon that could affect model parametrization was an apparent “self-inhibition” that occurred for certain species such that OD decreased after reaching its maximum value (e.g., *Clostridium symbiosum* in Fig. 5B). This phenomenon was rare in spent media and co-cultures, suggesting that it is relatively unimportant in community dynamics. In any case, the effects of self-inhibition on model parametrization, community dynamics, or both were evidently small relative to the effects of resource competition. Other interactions, such as the *Efe*-*Bt* interaction, persisted in larger communities. In these scenarios, we demonstrated that our framework could disentangle the mechanisms involved and incorporate the additional mechanisms into the model to improve predictions. We envision that this strategy can be executed iteratively to quantify other factors that contribute to community dynamics, such as interspecies differences in the efficiency of biomass generation and interspecies killing. In this manner, our framework provides a generalizable tool to construct mechanistic models of community dynamics for diverse communities in complex environments, which will facilitate the rational engineering of microbial communities.

## METHODS

### Bacterial culturing

Isolates were obtained via plating of fecal samples from humanized mice and frozen as glycerol stocks, as previously described (Aranda-Díaz *et al*., 2022). Frozen stocks were streaked onto BHI-blood agar plates (5% defibrinated horse blood in 1.5% w/v agar). Resulting colonies were inoculated into 3 ml Brain Heart Infusion (BHI) (BD #2237500) or modified Gifu Anaerobic Medium (mGAM) (HyServe #05433) in test tubes. All culturing and measurements were performed at 37 °C without shaking in an anaerobic chamber (Coy). To minimize potential physiological changes from freeze-thaw cycles and changes in growth medium, cultures were diluted 1:200 every 48 h for 3 passages before growth or metabolomics measurements. After the first passage, subsequent passages were performed in 96-well polystyrene plates (Greiner Bio-One) filled with 200 μl of growth medium.

### Bacterial growth measurements

Biomass yield over time was obtained via optical density at 600 nm (OD) as measured by an Epoch 2 plate reader (Biotek). All measurements were performed in clear, flat-bottomed 96-well plates (Greiner Bio-One #655161). Each well was filled with 200 μl of growth medium and inoculated with 1 μl of stationary phase culture immediately before measurement. Plates were sealed with transparent seals (Excel Scientific #STR-SEAL-PLT), with small (∼0.5 mm) holes cut above each well to allow gas exchange. Measurements were taken with continuous shaking at 37 °C.

### Growth in spent media

Spent media were obtained by centrifuging saturated cultures at 4,000 × 𝑔 for 5 min and filtering the supernatant with 0.22 μm polyethersulfone filters (Millex-GP #SLGP033RS) or 96-well 0.22 μm filter plates (Pall #8019). To investigate pH-mediated effects, *Bp*-spent medium was adjusted to a pH of 7.35 with NaOH, and filtered again to sterilize.

### Liquid chromatography-mass spectrometry (LC-MS) metabolomics

Spent media were collected as described above, and immediately stored at −80 °C. Samples were thawed only once immediately before LC-MS/MS analysis. Samples were analyzed by two chromatography methods, reversed phase (C18) and hydrophilic interaction chromatography (HILIC). Protocol details and parameters are described in the Supplemental Information. Briefly, metabolites were extracted using extraction mixtures containing stable isotope labeled internal standards. Samples for C18 analysis were dried at room temperature using a Labconco CentriVap, and reconstituted in 20% acetonitrile prior to analysis. 2 μl of prepared samples were injected onto a Waters Acquity UPLC BEH Amide column with an additional Waters Acquity VanGuard BEH Amide pre-column (HILIC) or Agilent SB-C18 column with a Phenomenex KrudKatcher Ultra filter frit attached to the column inlet (C18). The columns were coupled to a Thermo Vanquish UPLC machine. Chromatographic separation parameters (Showalter et al., 2018) and mass spectral parameters (Han *et al*., 2021) were described previously, with minor modifications (Supplemental Information). Spectra were collected using a Thermo Q Exactive HF Hybrid Quadrupole-Orbitrap mass spectrometer in both positive and negative mode ionization (separate injections, sequentially). Full MS-ddMS2 data were collected. Data were processed using MS-DIAL v. 4.60 (Tsugawa et al., 2015; Tsugawa et al., 2020). Alignment retention time and mass tolerance were set to 0.05 min and 0.015 Da, respectively. Aligned peaks were retained for further analyses only if they were present in at least two of three replicates and were >5-fold higher than the water blank average in at least one sample.

### Assembly experiments

Communities were assembled from stationary phase cultures of isolates mixed at equal volume, and 1 μl of the mixture was inoculated into 200 μl of growth medium. Plates were sealed and incubated at 37 °C without shaking. The assemblies were diluted 1:200 into fresh medium every 48 h for 5 passages to reach an ecological steady state in which further passages have virtually identical dynamics (Aranda-Díaz *et al*., 2022). The 15 single species “dropout” assemblies with 14 of the 15 members were passaged for 3 passages. In “refill” experiments, the inoculum for each dropout was mixed 1:1, 1:10, 1:100, 1:1,000, or 1:10,000 with the monoculture of the species that was left out, and passaged 3 times. The final passage for assembly experiments was grown in a plate reader for OD measurements, after which the plate was stored at −80 °C until DNA extraction for 16S rRNA gene sequencing was performed.

### 16S rRNA gene sequencing and analyses

Amplicon sequencing data were obtained and processed as previously described (Aranda-Díaz *et al*., 2022). Relative abundances were determined to a minimum threshold of 10^-4^, reflecting the typical depth of sequencing. The relative abundances of undetected species were set to 10^-4^ for visualization and for calculating the error between model predictions and experimental data. The three *Enterococcus* species were indistinguishable by the amplicon protocol used here. When more than one was present, their relative abundances were summed and visualized as *Eh* if *Eh* was present, else as *Efs*.

### Analyses of growth curves

Each growth curve in monoculture was fit to Eq. 2 with one resource to extract the final yield 𝐾, growth rate 𝜆, and lag time 𝜏 associated with that species. The culture yield over time 𝑋(𝑡) in Eq. 2 with one resource reduces to 𝑋(𝑡) = 𝐾[1 + (𝐾/𝑋_0_ − 1) exp(−𝜆(𝑡 − 𝜏))]^−1^, where 𝑋_0_ ≔ 𝑋(𝑡 = 0) is the initial value, and the final yield is taken at 48 h, 𝐾 ≔ 𝑋(𝑡 = 48 h). The growth rate and lag time were determined by a grid search to find the values that minimize the mean squared error between predicted and experimentally measured 𝑋(𝑡).

### Analyses of metabolomics data

Metabolomic features that passed pre-processing were defined as depleted or produced if they decreased by >100-fold or increased by >10-fold, respectively, compared to fresh medium, and if the difference was significant (*p*<0.05) by a two-sample *t*-test. Coarse-grained resources were obtained by grouping metabolomic features that shared the same set of consuming species.

### Simulations of coarse-grained consumer-resource (CR) model

To mimic our experimental protocol, Eq. 2 was simulated under a serial dilution scheme in which each dilution cycle continued until stationary phase when all resources were depleted (𝑑𝑌_𝜇_/𝑑𝑡 = 0 for all 𝜇), after which a new cycle was initiated by replenishing the resources to their initial levels 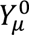 and diluting all species abundances by a factor 𝐷, which was set at 200 both experimentally and in simulations throughout this work. In simulations, the first cycle was initialized with equal abundances of each species, and dilutions were repeated until an ecological steady state was reached in which further cycles produced identical dynamics. At ecological steady state, species abundances in stationary phase are linear combinations of the resource levels since all resources have been converted to biomass. To compare against experimental data, relative abundances less than 10^-4^ were considered undetectable and removed in further calculations.

### Residues in randomly generated coarse-grained CR models

To determine the typical distribution of resource competition residues in coarse-grained CR models, we randomly selected 100 coarse-grained resources out of the 2^15^ − 1 possible groupings of 15 species. Each resource was assigned a random level from a uniform distribution from 0 to 1. The yields of monocultures and pairwise spent media experiments can then be calculated directly by summing the levels of resources consumed. Simulated yields were then modified with a 5% noise, the typical standard deviation of the mean in our measurements of yield, before calculating the resource competition residues.

### Parametrization of coarse-grained CR models

The parameters of the CR model in Eq. 2 are the initial resource levels 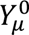 and resource consumption rates 𝑅_𝑖𝜇_ . The resource levels were inferred via linear regression as described in the text. Each experiment in monoculture and pairwise spent media represented one equation in the regression. In each equation, the unknowns are the resource levels that sum to the known final yield in that experiment. The consumption rates were inferred as described in the text.

Regularized linear regression, i.e., LASSO, was used to parametrize resource utilization structures with more unknowns than experiments. The regression problem was set up as in the linear regression case, with the addition that a regularization parameter determined the predicted number of coarse-grained resources with non-zero resource levels. The regularization parameter was chosen so that LASSO resulted in 40 non-zero resources.

### Parametrization of a generalized Lotka-Volterra (gLV) model

A gLV model was considered in which 𝑑𝑋_𝑖_/𝑑𝑡 = 𝑋_𝑖_(𝑟_𝑖_ + ∑_𝑗_ 𝐴_𝑖𝑗_𝑋_𝑗_), where 𝑋_𝑖_ denotes the abundance of species 𝑖, 𝑟_𝑖_ its growth rate, and 𝐴_𝑖𝑗_ the interaction coefficient of species 𝑗 on species 𝑖 . Model parameters 𝑟_𝑖_ and 𝐴_𝑖𝑗_ were parametrized from growth measurements in isolate and pairwise spent media as follows. In the absence of other species, species 𝑖 will reach a steady state abundance 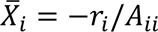. The self-interaction terms 𝐴_𝑖𝑖_ are free parameters that we set to −1 for simplicity. Hence, the growth rates are simply equal to the experimentally determined isolate yields, i.e., 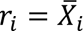. Then for the case of species 𝑗 growing in the spent medium of species 𝑖, we assumed that the effect of the spent medium was implemented within the model by a constant presence of species 𝑖 at its steady state value 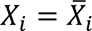. Thus, at steady state, the abundance of species 𝑗 grown in the spent medium of species 𝑖 is 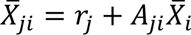. The interaction coefficient can therefore be expressed as a combination of experimentally determined yields in spent media, i.e., 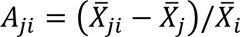.

## ACKNOWLEDGMENTS

We thank members of the Huang lab for helpful discussions, and Jonas Cremer, Ben Good, Karna Gowda, Mikhail Tikhonov, Ned Wingreen, and Katherine Xue for a critical reading of the manuscript. We thank Biohub team member Wasim Sandhu for metabolite extraction and LC-MS/MS data acquisition. This work was funded by the Stanford School of Medicine Dean’s Postdoctoral Fellowship (to P.H.), NIH Postdoctoral Fellowship F32 GM143859 (to P.H.), NSF Graduate Research Fellowship (to T.H.N.), NSF Awards EF-2125383 and IOS-2032985 (to K.C.H.), and NIH Awards R01 AI147023 and RM1 GM135102 (to K.C.H.). K.C.H. is a Chan Zuckerberg Biohub Investigator.

## SUPPLEMENTAL FIGURES

**Figure S1:**
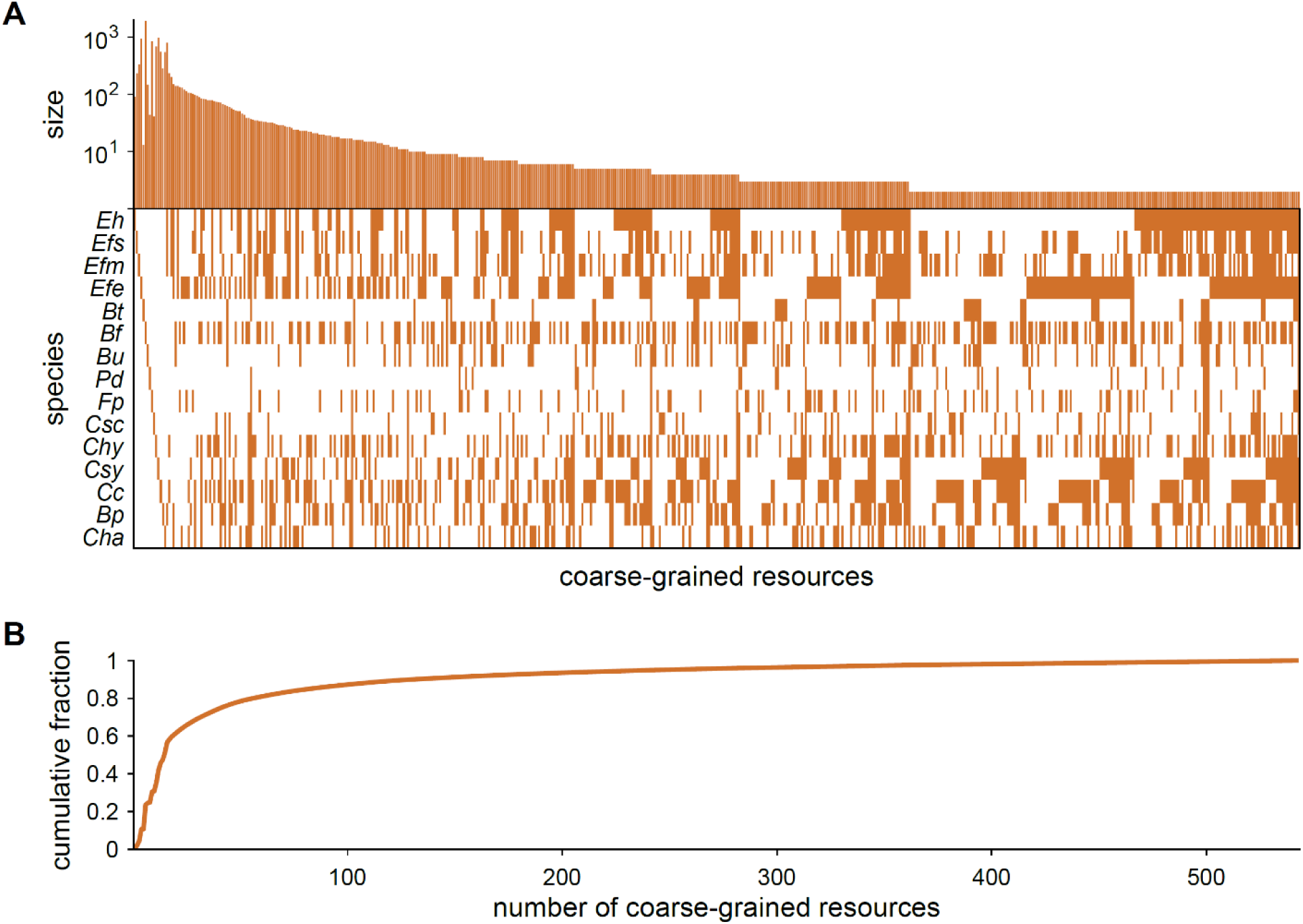
Metabolomics-derived coarse-grained resources. A) Size and structure of metabolomics-derived coarse-grained resources. A metabolomic peak was considered depleted if it decreased by >100-fold compared to fresh media. Metabolites that share the same set of consuming species were grouped together and are shown as one column in the matrix. The number of metabolomic peaks in each group is shown above each column. Only niches with more than one constituent metabolite are shown. B) The cumulative fraction of the number of metabolomic peaks as a function of the number of groups.

**Figure S2:**
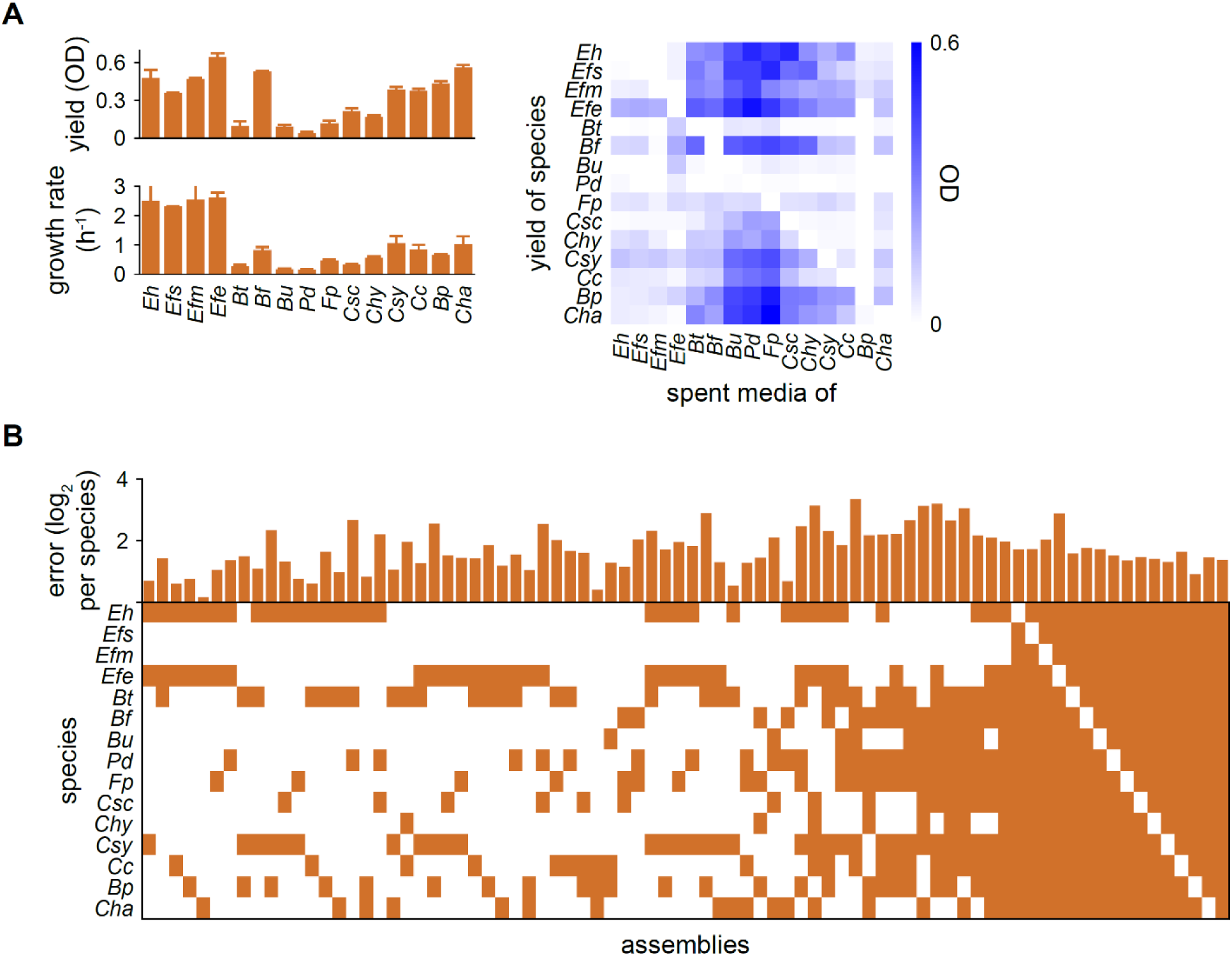
CR model successfully predicted assembly compositions. A) Model input: growth rates and yields in monocultures (left) and yields in pairwise spent media (right). The mean value across 2-4 replicates is shown. Error bars denote the standard error of the mean. Parametrization outputs are the inferred resource levels and consumption rates, and are shown in Fig. 3B. B) Prediction error for each assembly. Only assemblies with more than 2 species are shown. All pairwise co-cultures were also assembled and tested. For each assembly, the error was calculated between model predictions and the mean relative abundance observed across 3 experimental replicates.

**Figure S3:**
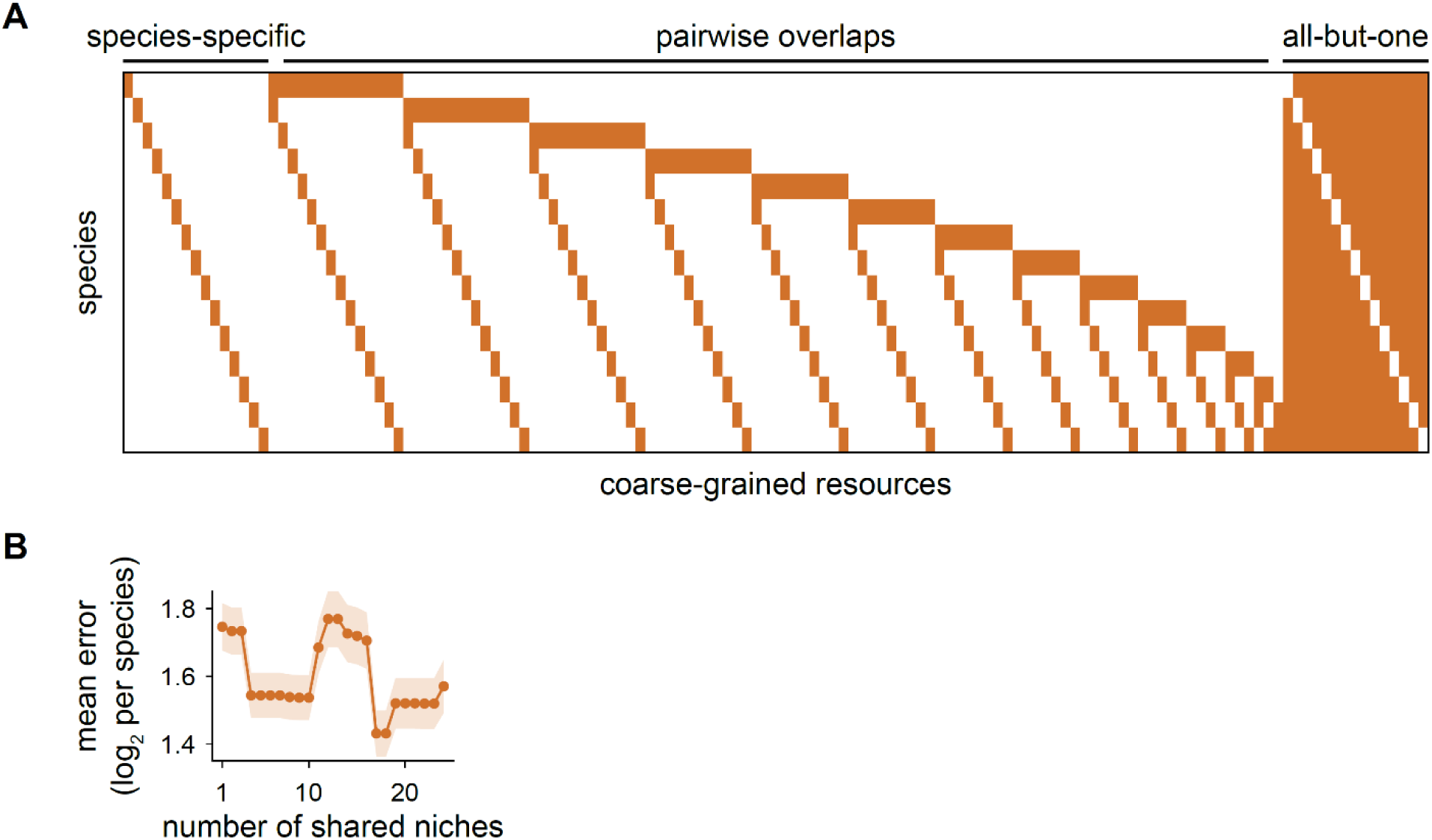
Hypothetical structures of resource utilization failed to accurately predict assembly compositions. A) Hypothetical structures of resource utilization. The species-specific niches are consumed by only one species, and form the set of niches in the “base” structure. The pairwise overlaps are consumed by only two species. The all-but-one niches are consumed by 14 of the 15 species. Model performance using the base structure, the base structure with the pairwise overlaps, and the base structure with the set of all-but-one resources are shown in Fig. 3C. B) Mean errors for the base structure plus a varying number of the largest remaining niches are shown. Shaded region denotes standard error of the mean.

**Fig S4:**
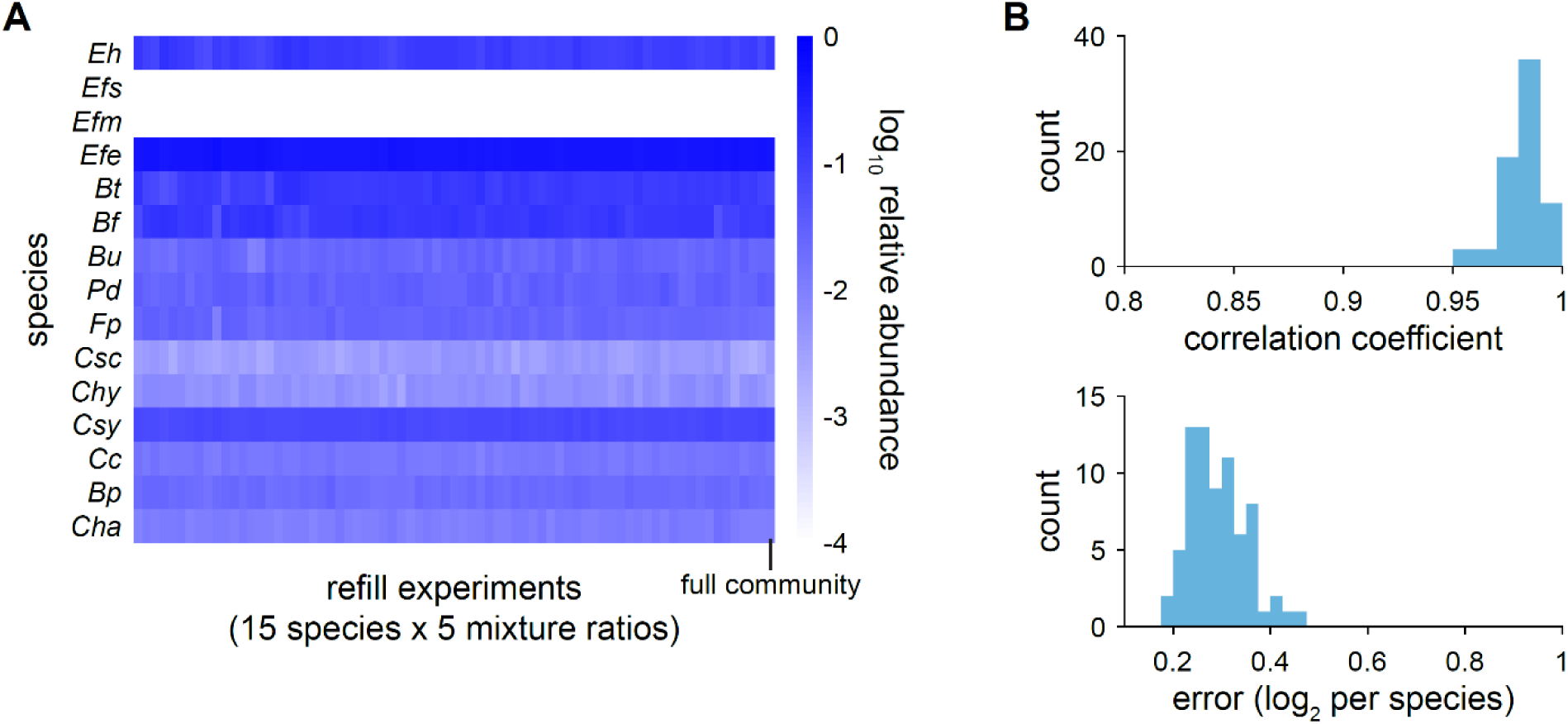
Assembly compositions were independent of initial values. A) Relative abundances in “refilled” dropout assemblies. Each column represents one experiment, in which a dropout assembly with 14 of the 15 species was mixed with the monoculture of the species that was left out, at varying ratios (1:1, 1:10, 1:100, 1:1,000, and 1:10,000). All conditions (15 species × 5 ratios) are shown except for 3 experiments with idiosyncratic sequencing errors. The compositions were virtually indistinguishable from each other and from the full 15 member community, which is shown in the last column. B) Histogram of the correlation coefficient (top) and error per species (bottom) between the species compositions in each refill experiment and the full 15 member community.

**Figure S5:**
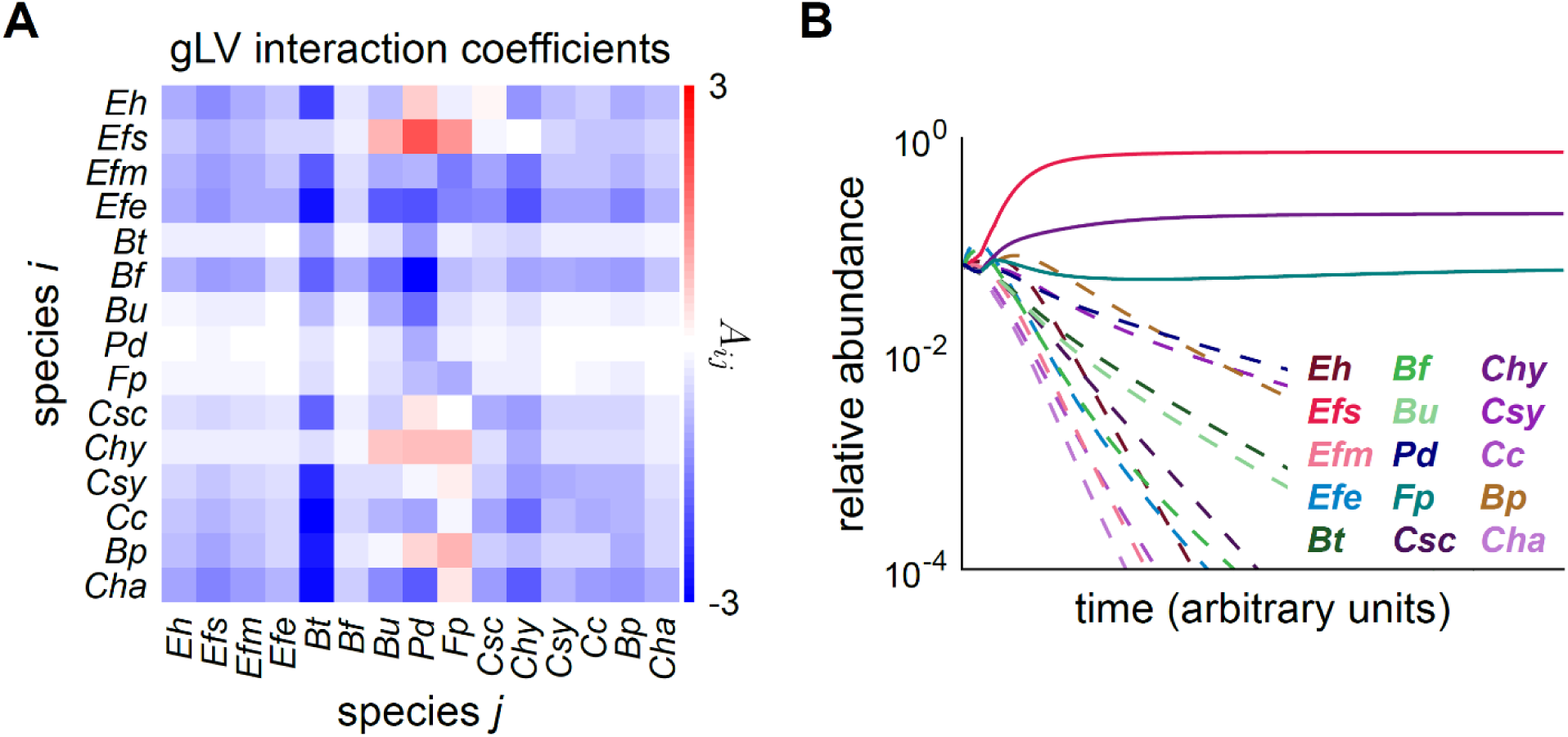
A generalized Lotka-Volterra (gLV) model failed to accurately predict assembly compositions. A) The matrix of interaction coefficients inferred for growth in BHI is shown (Methods). B) Model predictions for the dynamics of the community with all 15 species are shown. Only 3 species coexisted at steady state, in stark disagreement with experiment.

**Figure S6:**
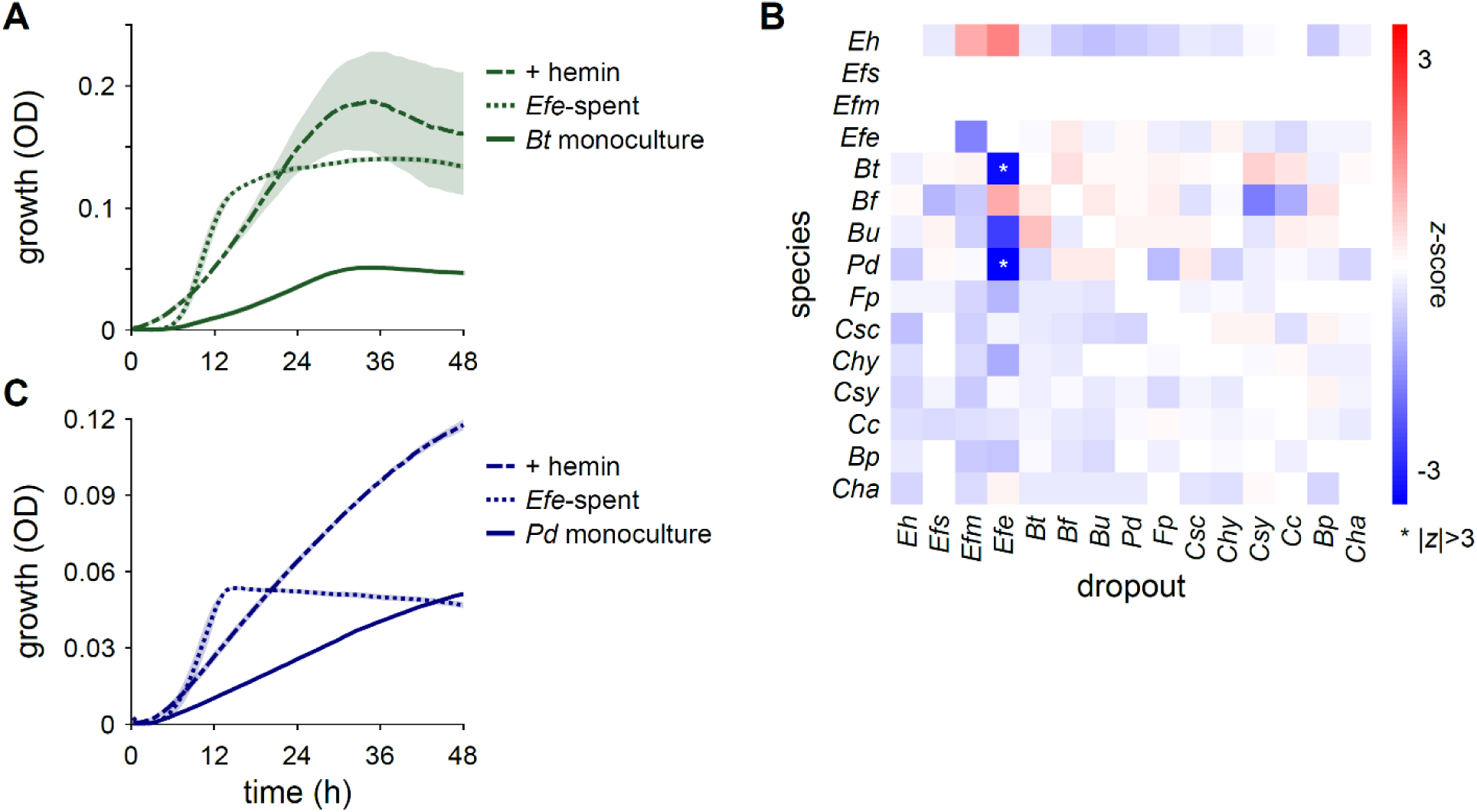
Strong interactions in dropout assemblies were rare. A) Optical density (OD) over time for *Bt* grown in monoculture (solid line), in *Efe*-spent medium (dotted line), and in fresh BHI plus hemin (dash dotted line). The mean over 2-3 replicates is shown, and shading denotes standard error of the mean. B) *z-*scores in dropout assemblies. Each column represents a dropout assembly of 14 of the 15 species, with the denoted species left out of the community. Each row represents the *z*-scores calculated from the relative abundances of the denoted species. *z*-scores are defined as 𝑧_𝑖𝑗_ ≔ (𝑥_𝑖𝑗_ − 𝜇_𝑖_)/𝜎_𝑖_, where 𝑥_𝑖𝑗_ is the log_10_ relative abundance of species 𝑖 in the dropout assembly in which species 𝑗 was left out, and 𝜇_𝑖_and 𝜎_𝑖_are the mean and standard deviation, respectively, of the log_10_ relative abundance of species 𝑖 across all dropout assemblies. *z*-scores with absolute value larger than 3 are denoted by an asterisk. C) Same as (a) but for *Pd*.

